# TRPV4 stimulates colonic afferents through mucosal release of ATP and glutamate

**DOI:** 10.1101/2024.04.12.589184

**Authors:** Michelle Y Meng, Luke W Paine, David Sagnat, Ivana Bello, Sophie Oldroyd, Farideh Javid, Matthew T Harper, James RF Hockley, Ewan St. J Smith, Róisín M Owens, Laurent Alric, Etienne Buscail, Fraser Welsh, Nathalie Vergnolle, David C Bulmer

**Affiliations:** Department of Pharmacology, University of Cambridge, Cambridge, UK; IRSD, Université de Toulouse, INSERM, INRAE, ENVT, Univ Toulouse III-Paul Sabatier (UPS), Toulouse, France; Department of Pharmacy, School of Medicine and Surgery, University of Naples Federico II, 80131 Naples, Italy; Department of Chemical Engineering and Biotechnology, University of Cambridge, Cambridge, UK; Department of Pharmacy, School of Applied Sciences, University of Huddersfield, Huddersfield, UK; Internal Medicine Department of Digestive Disease CHU Toulouse-Rangueil and Toulouse University, UPS, 31000 Toulouse, France; Department of Surgery, CHU Toulouse-Rangueil and Toulouse University, UPS, 31059 Toulouse, France; Biopharmaceuticals R&D, AstraZeneca, Neuroscience, Cambridge, UK; Department of Physiology & Pharmacology, University of Calgary Cumming School of Medicine, 3330 Hospital Drive NW, Calgary, Ab T2N 4N1, Canada

**Keywords:** TRPV4, colonic afferents, adenosine triphosphate, glutamate, intestinal mucosa, visceral hypersensitivity

## Abstract

**Background and Purpose:** Abdominal pain is a leading cause of morbidity for people living with gastrointestinal disease. While the vanilloid transient receptor potential 4 (TRPV4) ion channel has been implicated in the pathogenesis of abdominal pain, the relative paucity of TRPV4 expression in colon-projecting sensory neurons suggests that non-neuronal cells may also contribute to TRPV4-mediated nociceptor stimulation.

**Experimental Approach:** Changes in murine colonic afferent activity were examined using *ex vivo* electrophysiology in tissues with the gut mucosa present or removed. ATP and glutamate release were measured by bioluminescence assay from human colon organoid cultures and mouse colon. Dorsal root ganglion sensory neuron activity was evaluated by Ca^2+^ imaging when cultured alone or co-cultured with colonic mucosal cells.

**Key Results:** The TRPV4 agonist GSK1016790A elicited a robust increase in murine colonic afferent activity, which was abolished by removal of the gut mucosa. GSK1016790A promoted ATP and glutamate release from human colon organoid cultures and mouse colon. Inhibition of ATP degradation in mouse colon enhanced the afferent response to GSK1016790A. Pre-treatment with purinoreceptor or glutamate receptor antagonists attenuated and abolished the response to GSK1016790A when given alone or in combination, respectively. Sensory neurons co-cultured with colonic mucosal cells produced a marked increase in intracellular Ca^2+^ to GSK1016790A compared to neurons cultured alone.

**Conclusions and Implications:** Our data indicate that mucosal release of ATP and glutamate is responsible for the stimulation of colonic afferents following TRPV4 activation. These findings highlight an opportunity to target the gut mucosa for the development of new visceral analgesics.

**Bullet Point Summary:** *What is already known?*

- Activation of TRPV4 causes visceral hypersensitivity via the stimulation of colonic afferents.

*What does this study add?*

- TRPV4-mediated colonic afferent activation is dependent on mucosal release of ATP and glutamate.

*What is the clinical significance?*

- Mucosal TRPV4-mediated colonic afferent activation provides a gut restricted target for treating abdominal pain.

## 1. Introduction

The vanilloid transient receptor potential 4 (TRPV4) channel is a non-selective cation channel which displays greater conductance for divalent over monovalent ions resulting in the influx of extracellular Ca^2+^ upon activation. TRPV4 is a polymodal sensor of heat and hypo-osmolality and can be activated by endogenous lipids including arachidonic acid and its epoxyeicosatrienoic acid (EET) metabolite 5,6-EET (Güler et al., 2002; Liedtke et al., 2000; Watanabe et al., 2003). While TRPV4 appears to play roles in homeostatic function, such as detecting extracellular fluid osmolality and shear stress-induced vasodilation, it has also been implicated in disease (Köhler et al., 2006; Lechner et al., 2011). For example, levels of 5,6-EET are elevated in colonic biopsies from patients with diarrhoea-predominant irritable bowel syndrome (IBS-D) and correlate with the magnitude and frequency of abdominal pain, thus linking TRPV4 activation to disease pathophysiology in IBS-D (Cenac et al., 2015). Consistent with these findings, stimulation of TRPV4 with 5,6-EET and other agonists such as 4α-phorbol 12, 13-didecanoate (4α-PDD) or GSK1016790A is pro-nociceptive, increasing intracellular [Ca^2+^] in sensory neurons, promoting nerve discharge in afferent fibres innervating the bowel, and enhancing pseudoeffective pain behaviours such as the visceromotor response (VMR) evoked by colorectal distension (CRD) (Brierley et al., 2008; Cenac et al., 2008; McGuire et al., 2018; Poole et al., 2013; Sipe et al., 2008). In addition, TRPV4 serves as a downstream effector of visceral hypersensitivity evoked by other algogenic and inflammatory mediators elevated in the bowel of IBS-D patients such as tryptase, serotonin, and histamine. These effects are driven by the potentiation of TRPV4 currents due to increased membrane expression and phosphorylation of tyrosine and serine/threonine residues in TRPV4 following cognate receptor activation i.e. protease-activated receptor 2 (PAR2) or histamine H_1_ receptors (Cenac et al., 2010; Fan et al., 2009; Poole et al., 2013). In particular, TRPV4 is critical for the activation of colonic afferents and development of visceral hypersensitivity evoked by PAR2 due to the generation of 5,6-EET following PAR2 stimulation (Cenac et al., 2008; Grant et al., 2007; Poole et al., 2013; Sipe et al., 2008).

Our recent single cell RNA sequencing (scRNA-seq) analysis of colonic sensory neurons revealed that TRPV4 mRNA is present in a relatively small population of colonic neurons (Hockley et al., 2019). This is in contrast to the marked excitatory effect of TRPV4 on colonic afferent activity discussed above and thus suggests that non-neuronal cell types may provide a substantial contribution to TRPV4-mediated colonic afferent responses by releasing secondary signalling mediators. Consistent with this hypothesis, TRPV4-mediated ATP release has previously been demonstrated from the oesophageal epithelium and in astrocytes where TRPV4 also evokes glutamate release (Mihara et al., 2011; Shibasaki et al., 2014). These observations prompted us to evaluate the contribution of ATP and glutamate release from the gut in TRPV4-mediated colonic afferent activation.

## 2. Methods

### 2.1 Materials

Stock concentrations were prepared for all drugs as recommended and diluted to working concentrations on the day of testing. GSK1016790A, ATP, D-aminophosphonovaleric acid (D-AP5), and capsaicin were purchased from Sigma Aldrich. HC067047, CGS 15943, pyridoxalphosphate-6-azophenyl-2’,4’-disulfonic acid (PPADS), sodium polyoxotungstate (POM-1), bradykinin, ASP7663, and 2,3-dihydroxy-6-nitro-7-sulfamoylbenzo(f)quinoxaline (NBQX) were purchased from Tocris. 3-[(2-methyl-1,3-thiazol-4-yl)ethynyl]pyridine (MTEP) hydrochloride was purchased from Abcam, and RO4 was a gift from Dr Wendy Winchester.

### 2.2 Ethical approval

All animal experiments were conducted in accordance with the Animals (Scientific Procedures) Act 1986 Amendment Regulations 2012 and following local ethical review by the University of Cambridge Animal Welfare and Ethical Review Body (AWERB).

### 2.3 Animals

Experiments were performed using tissue from male C57BL/6J mice (8-14 weeks) purchased from Charles River (Cambridge, UK, RRID:IMSR_JAX:000664). This species and strain were selected to allow us to compare our findings with previous studies exploring investigating TRPV4-associated roles in the gut and visceral hypersensitivity (Cenac et al., 2015; Sipe et al., 2008). Mice were conventionally housed in groups of up to 8 per cage with nesting material, enrichment (shelter, tubes, and chewing blocks), and access to food and water *ad libitum*. Rooms were temperature-controlled (21 °C) with a 12-hour light/dark cycle. Animals were euthanised by a rising concentration of CO_2_ followed by exsanguination.

### 2.4 Ex vivo electrophysiological recording from the lumbar splanchnic nerve

#### Nerve Recording

Electrophysiological recordings were conducted as previously described (Barker et al., 2023). Briefly, following euthanasia, the colorectum with associated lumbar splanchnic nerves (LSN) was isolated. Tissue was cannulated as a tubular preparation in a recording bath with a Sylgard base (Dow Corning, UK), luminally perfused (200 µL min^-1^) by a syringe pump (Harvard Apparatus, MA), and serosally superfused (7.5 mL min^-1^, 31-34 °C) with carbogenated (95% O_2_-5% CO_2_) Krebs buffer (in mM: NaCl 124, KCl 4.8, NaH_2_PO_4_ 1.3, CaCl_2_ 2.5, MgSO_4_·7H_2_O 1.2, D(+)-glucose 11.1, NaHCO_3_ 25). Krebs buffer was supplemented with atropine (10 µM, Sigma Aldrich) and nifedipine (10 µM, Sigma Aldrich) to inhibit smooth muscle activity (Bhebhe et al., 2023; Cibert-Goton et al., 2021). For mucosa-free preparations, the tissue was opened along the mesenteric border, pinned with the mucosal side up and the mucosal layer gently dissected away (Figure S1a-d). Complete removal of the mucosa was confirmed using hematoxylin and eosin staining (Figure S1e,f). Confirmation of spontaneous contractility and comparison of baseline activity before and after mucosal removal were used to ensure tissue viability was retained (Figure S1g). Multi-unit LSN activity was recorded from teased nerve bundles using borosilicate glass suction electrodes. Signals were amplified (gain 5 kHz), bandpass filtered (100-1300 Hz; Neurolog, Digitimer Ltd, UK), and digitally filtered (Humbug, Quest Scientific, Canada) to remove 50 Hz noise. Data were digitised at 20 kHz (micro1401; Cambridge Electronic Design, UK) prior to display using Spike 2 software (Cambridge Electronic Design, UK).

#### Electrophysiology Protocols

Data were collected following a 30-minute stabilisation period to establish baseline firing with a signal-to-noise ratio sufficient to allow accurate spike counting. Agonists and antagonists were diluted from stock solutions into final volumes of 20 mL or 50 mL Krebs respectively and applied by serosal superfusion via the in-line heater. A washout period of 30-minutes was given between repeat drug applications on the same tissue preparation. Antagonist studies were conducted blinded except for coloured drug PPADS (red/orange) where blinding was not possible.

#### Data analysis

Action potentials were determined by counting spiking waveforms passing through a threshold set at twice the background noise (typically 30-50 µV) and ongoing nerve discharge was expressed as a mean frequency (time base 60-seconds). The change in nerve activity post-treatment was determined by subtracting a baseline pre-treatment firing rate (mean firing for the 6-minute period prior to treatment) from the mean firing rate. Data from individual experiments were averaged to give a mean and standard error of the mean (SEM) value of nerve discharge over time. The area under the curve (AUC) was calculated as an indicator of multiunit nerve activity for 900-seconds following initial GSK1016790A, bradykinin, or ATP application.

### 2.5 Primary culture of mouse dorsal root ganglion neurons

Dorsal root ganglion (DRG) neurons were cultured as previously described (Higham et al., 2024). Briefly, isolated DRG (T12-L5, spinal segments innervating the distal colon) were incubated in 1 mg mL^-1^ type 1A collagenase (15-minutes) followed by 1 mg mL^-1^ trypsin (30-minutes) both with 6 mg mL^-1^ BSA in Leibovitz’s L-15 Medium, GlutaMAX™ Supplement (supplemented with 2.6% (v/v) NaHCO_3_). DRG were resuspended in 2 mL Leibovitz’s L-15 Medium, GlutaMAX™ Supplement containing 10% (v/v) FBS, 2.6% (v/v) NaHCO_3_, 1.5% (v/v) D(+)-glucose, and 300 units mL^-1^ penicillin and 0.3 mg mL^-1^ streptomycin (P/S). DRG were mechanically dissociated, centrifuged (100 *g*), and the supernatant collected for five triturations. Following centrifugation and resuspension, the supernatant (50 µL) was plated onto poly-D-lysine-coated 35 mm glass bottom culture dishes (MatTek, MA, USA), further coated with laminin (Thermo Fisher). Dishes were incubated for 3-hours to allow cell adhesion, before flooding with 2 mL supplemented L-15 medium and cultured for 24-hours. All incubations were carried out at 37 °C with 5% CO_2_.

### 2.6 Primary culture and co-culture of mouse colonic mucosal cells

The protocol for mucosal cell culture was adapted from Psichas et al., 2017. Luminal contents were flushed from the colon with cold PBS and the outer muscle layer removed. Tissue was minced and digested with 0.35 mg mL^-1^ type 1A collagenase (5x 10-minutes). Crypt containing fractions were centrifuged (100 *g*) for 3-minutes and resuspended with DMEM supplemented with 10% (v/v) FBS, 2% (v/v) NaHCO_3_, 1.5% (v/v) glucose, P/S, and Y-27632 (10 μM), before filtering through a 100 μm cell strainer. Crypts were plated at a density of 1000 crypts per cm^2^ onto poly-D-lysine-coated 35 mm glass bottom culture dishes (MatTek, MA, USA), further coated with laminin (Thermo Fisher). Dishes were incubated for 3-hours before flooding with 2 mL supplemented DMEM. For co-culture experiments, DRG were isolated and cultured as described then plated on top of mixed colonic mucosal cell cultures isolated 24-hours previously and flooded before use 24-hours later. All incubations were carried out at 37 °C with 5% CO_2._

### 2.7 RNA extraction and polymerase chain reaction

Total RNA was isolated from primary cultured colonic mucosal cells using the RNeasy® Mini Kit and QIAshredder (Qiagen) according to the manufacturer’s instructions and DNase (Promega) treated to maximise RNA quality. 500 ng RNA was subjected to reverse transcription (RT) to cDNA using the High-Capacity cDNA Reverse Transcription Kit (Thermo Fisher). A negative RT reaction was performed without reverse transcriptase to check for genomic DNA contamination during the PCR. PCR was performed with specific primer sets (Merck) directed against TRPV4 and β-actin (Table 1) with cycling conditions of: one cycle of 95 °C for 30-seconds, and 30 cycles of 95 °C for 30-seconds, 55 °C for 1-minute, and 72 °C for 1-minute.

**Table 1.**
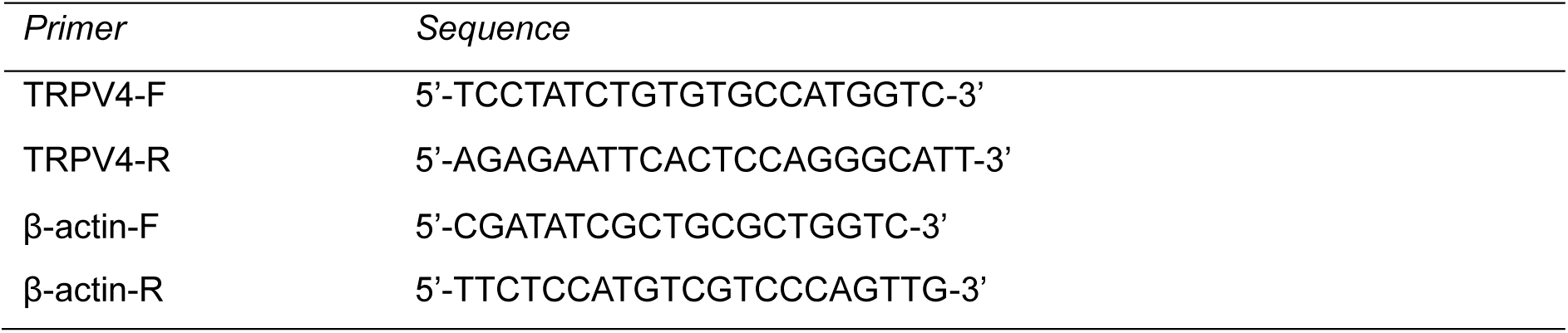
Primers designed for the amplification of TRPV4 and housekeeping gene β-actin based upon the reference genome for mouse (*Mus musculu*s, GRCm39).

### 2.8 Colon Histology and Staining

Mucosal tissue removed from the colon and the underlying muscle was fixed in 4% (w/v) paraformaldehyde (Sigma Aldrich) overnight at 4 °C followed by cryopreservation in 30% (w/v) sucrose overnight (Sigma Aldrich) at 4 °C. Tissue was embedded in M-1 Embedding Matrix (Thermo Fisher Scientific) and snap frozen in liquid nitrogen. Embedded tissue was sectioned into 10 μm slices using a Leica CM3000 cryostat and mounted on SuperFrost slides (Thermo Fisher Scientific). Sections were stained with hemotoxylin (0.2% w/v; Sigma Aldrich) and eosin (0.5% w/v; Acros Organics) to confirm mucosa removal. Slides were mounted with 70% (v/v) glycerol and imaged using an Olympus Bx51 microscope.

### 2.9 Immunocytochemistry

Unless otherwise stated, all incubations were carried out at room temperature. Antibodies used are summarised in Table 2. MatTek plated cells were fixed with 4% paraformaldehyde for 10-minutes, permeabilized with 0.05% TritonX-100 for 5-minutes, and blocked in 5% donkey serum in 0.2% Triton-X100 for 30-minutes. Cells were incubated in primary antibody for 24-hours at 4°C. Secondary antibodies were incubated for 1-hour followed by 4’-6-diamidino-2-phenylindole (DAPI; 1:1000, Abcam) for 5-minutes. Immunostained cells were imaged on an Olympus Bx51 microscope and images were captured on a Qicam CCD camera (Qimaging) with a 100-millisecond exposure and false coloured (βIII-tubulin, green; DAPI, blue; E-cadherin, vimentin; red). Staining was not observed when primary antibodies were omitted (data not shown).

**Table 2.**
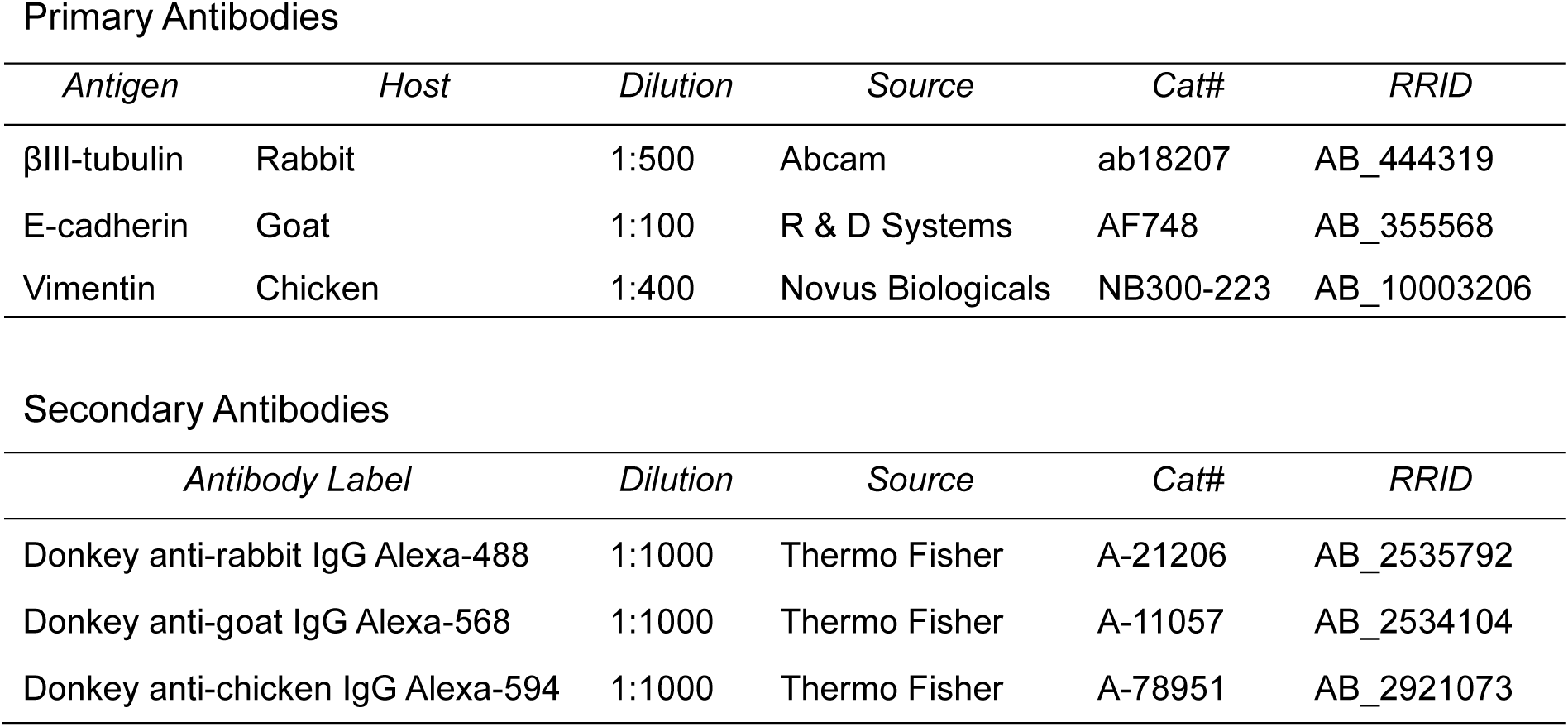
Antibodies use for immunohistochemistry of primary DRG and colonic mucosal cell co-cultures. *IgG; immunoglobulin G*.

Primary Antibodies
| *Antigen* | *Host* | *Dilution* | *Source* | *Cat#* | *RRID* |
| --- | --- | --- | --- | --- | --- |
| βIII-tubulin | Rabbit | 1:500 | Abcam | ab18207 | AB_444319 |
| E-cadherin | Goat | 1:100 | R & D Systems | AF748 | AB_355568 |
| Vimentin | Chicken | 1:400 | Novus Biologicals | NB300-223 | AB_10003206 |

| *Antibody Label* | *Dilution* | *Source* | *Cat#* | *RRID* |
| --- | --- | --- | --- | --- |
| Donkey anti-rabbit IgG Alexa-488 | 1:1000 | Thermo Fisher | A-21206 | AB_2535792 |
| Donkey anti-goat IgG Alexa-568 | 1:1000 | Thermo Fisher | A-11057 | AB_2534104 |
| Donkey anti-chicken IgG Alexa-594 | 1:1000 | Thermo Fisher | A-78951 | AB_2921073 |

### 2.10 Ca^2+^ imaging

#### Data acquisition

Extracellular solution (ECS, in mM: NaCl 140, KCl 4, MgCl_2_ 1, CaCl_2_ 2, D(+)-glucose 4, HEPES 10) was prepared and adjusted to pH 7.4 using NaOH and an osmolality of 290-310 mOsm using sucrose. Cells were incubated for 30-minutes with 100 µl of 10 μM Ca^2+^ indicator Fluo-4-AM (37 °C; shielded from light). For inhibitor studies requiring pre-incubation, 200 µL of antagonist was added for 10-minutes prior to imaging. Dishes were mounted on the stage of an inverted microscope (Nikon Eclipse TE-2000S) and cells were visualised at 10 X magnification with brightfield illumination. Fluorescent images were captured with a CCD camera (Retiga Electro, Photometrics) at 2.5 fps with 100-millisecond exposure and a 470 nm filter cube (Cairn Research, Faversham, UK). Emission at 520 nm was recorded with μManager32. All protocols began with a 10-second baseline of ECS before drug superfusion to establish baseline fluorescence. Finally, cells were stimulated with 50 mM KCl for 10-seconds to determine cell viability, identify neuronal cells, and allow normalisation of fluorescence. A fresh dish was used for each protocol and all solutions were diluted in ECS.

#### Data analysis

Individual cells were circled on a brightfield image and outlines overlaid onto fluorescent images using ImageJ (NIH, MA, USA). Pixel intensity was measured and analysed with custom-written scripts in RStudio (RStudio, MA, USA). Background fluorescence was subtracted from each cell and fluorescence intensity (F) baseline corrected and normalised to the maximum fluorescence elicited during 50 mM KCl stimulation (F_max_). Maximum KCl fluorescence was denoted as 1 F/F_max_. Further analysis was confined to cells with a fluorescence increase ≥ 5 standard deviations away from the mean baseline before 50 mM KCl application. Neurons were deemed responsive to a drug challenge if a fluorescence increase of 0.1 F/F_max_ was seen in response to drug perfusion. The proportion of responsive neurons and magnitude of the fluorescence response were measured for each experiment, with peak responses calculated from averaging fluorescence values from individual neurons at each time point.

### 2.11 ATP and glutamate secretion assays

#### Mouse Colon Tissue Preparation

The ascending colon to rectum was removed, luminal contents flushed, and opened longitudinally. Incubations were performed at 37 °C in a 1.5 mL eppendorf tube containing Krebs solution with NTPDase 1-3 inhibitor POM-1 (100 μM). Samples were stabilised in 1 mL Krebs solution for 10-minutes before Krebs was replaced with GSK1016790A (0.1 - 10 μM) or DMSO (0.01%). For inhibition studies, samples were incubated with HC067047 (10 μM) or DMSO (0.01%) for 10-minutes prior to GSK1016790A (10 μM). After 10-minutes, supernatants were removed and cellular debris removed by centrifugation (200 *g*, 5-minutes, 4 °C).

#### Human Colon Organoid Preparation

Human tissue no longer required for pathological assessment was collected following informed patient consent from individuals undergoing surgical bowel resections as part of their standard clinical treatment for colorectal cancer or endometriosis at the Toulouse University Hospital (CODECOH national agreement DC2015-2443, COLIC Collection). Colon crypts were isolated and cultured as previously described (d’Aldebert et al., 2020). Organoid cultures were expanded and seeded on transwell inserts (24-well plates, 0.4 μm pore size, Corning COSTAR). Organoids were collected and dissociated into single cells by incubation in pre-warmed TrypLE Express Enzyme for 10-minutes at 37 °C in agitation (1000 rpm). After addition of 5 mL Advanced DMEM/F12 with 2 mM Glutamax, 10 mM HEPES, and 10% FBS, and centrifugation (400 rpm, 5-minutes, 4 °C), dissociated cells were resuspended in organoid culture medium (50:50 volume L-WRN CM and Advanced DMEM/F12, Glutamax/HEPES containing FBS (37.5:7 v/v DMEM/FBS). Cells were seeded at a density of 4 x 10^5^ on individual transwells pre-coated with Matrigel (1:40 v/v in PBS). 600 μL culture medium supplemented with SB431542 (10 μM) and Y-27632 (10 μM) was added to the basolateral side and cells were incubated at 37 °C in 5% CO_2_. Media was replaced every 2-3 days until cells reached confluence. On the day of the experiment, cells were washed with PBS, medium replaced with PBS in the basolateral compartment, and PBS containing GSK1016790A (1 μM) or DMSO (0.01%) was added to the apical compartment. Media from the basolateral compartment was collected before and 10-minutes after apical exposure to GSK1016790A or DMSO.

#### Secretion Assays

ATP secretion was determined using the CellTiter-Glo® 2.0 (Promega) ATP kit. Glutamate secretion was determined using Glutamate-Glo® Assay (Promega) with samples diluted in Krebs 1:40 prior to measurement. All samples were run as technical triplicates and luminescence measured with a plate reader (FluoStar Omega; BMG Labtech). Concentrations were calculated from luminescence signals using a standard curve generated from ATP or glutamate standards and linear regression using Prism 9 (GraphPad Software, USA).

### 2.12 Statistics

The data and statistical analyses in this study comply with the recommendations on experimental design and analysis in pharmacology (Curtis et al., 2022). Statistical analysis and compilation of figures was conducted using GraphPad Prism Version 9.0.0 for Windows (GraphPad, Inc.). Data were examined to ensure the assumptions for parametric analysis were fulfilled and the appropriate non-parametric analyses were used if required. Normality testing was performed using Shapiro-Wilk test. For two groups, Student’s *t* tests or Mann-Whitney U test were applied as appropriate. For pairwise comparison, paired Student’s *t* test was performed. Nerve response traces were analysing using two-way repeated-measures ANOVA. DRG and co-culture peak F/F_max_ values and % neuron responses were analysed using two-way ANOVA. For more than two groups, one-way ANOVA was used. Holm–Šídák’s multiple comparisons post-hoc test was performed if F achieved *P* < 0.05. Data are displayed as mean ± SEM. The level of probability deemed to constitute statistical significance was established as \**P* < 0.05.

## 3 Results

### 3.1 TRPV4-induced colonic afferent activation is inhibited by purinoceptor and NMDA receptor antagonists

Bath application of the selective TRPV4 agonist GSK1016790A (0.001-10 μM) evoked a concentration-dependent increase in colonic afferent firing which was abolished by pretreatment with the TRPV4 antagonist HC067047 (10 μM, *P* < 0.0001, Figure S2a,b) (Thorneloe et al., 2008; Willette et al., 2008). The response to GSK1016790A at 10 μM consisted of a robust, prolonged increase in nerve discharge which did not return to baseline within 30-minutes of treatment or elicit a further increase in firing to a second repeat application of GSK1016790A (Figure 1a, S2c).

**Figure 1.**
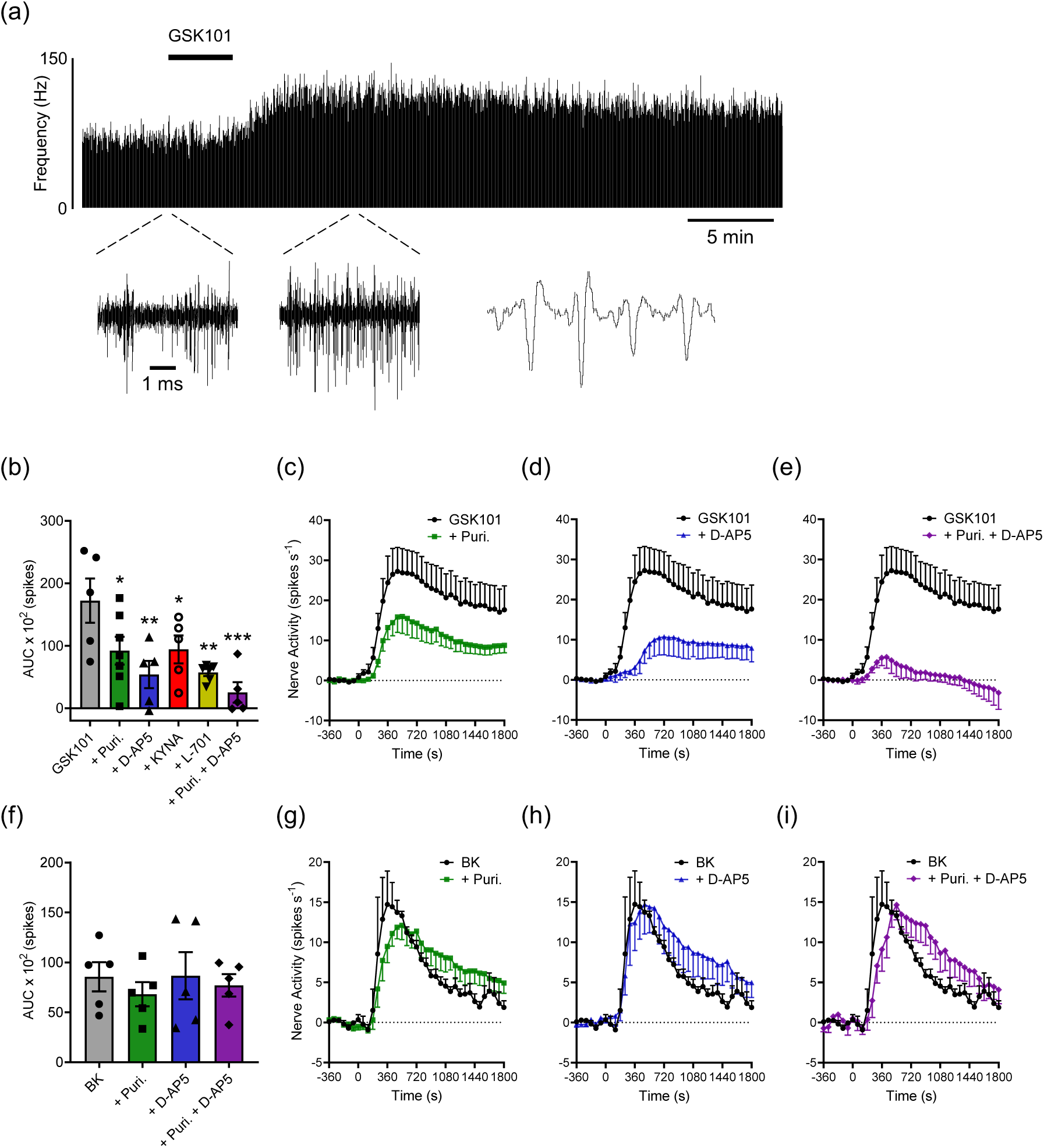
ATP and glutamate drive TRPV4-induced LSN firing. **(a)** Example rate histogram demonstrating an increase in LSN activity following application of GSK1016790A (GSK101, 10 μM). Bin width = 1-second. **(b)** AUC of nerve activity from t = 0-900 seconds following application of GSK1016790A (10 μM) alone or following pretreatment with the purinoceptor antagonist cocktail (Puri.: CGS 15943 (3 μM), RO4 (10 μM), and PPADS (100 μM)), D-AP5 (200 μM), kynurenic acid (KYNA, 30 μM), L-701,324 (L-701, 100 nM), or the purinoceptor antagonist cocktail and D-AP5 combined. One-way ANOVA with Holm–Šídák’s multiple comparisons test compared to GSK1016790A only group (*N* = 5-7). **(c)** Change in nerve activity following application of GSK1016790A (10 μM) alone or following pretreatment with the purinoceptor antagonist cocktail. **(d)** Change in nerve activity following application of GSK1016790A (10 μM) alone or following pretreatment with D-AP5. **(e)** Change in nerve activity following application of GSK1016790A (10 μM) alone or following pretreatment with the purinoceptor antagonist cocktail and D-AP5. GSK1016790A only dataset repeated in (c), (d), and (e) for ease of interpretation. (c-e) Two-way repeated-measures ANOVA (*N* = 5-7). **(f)** AUC of nerve activity from t = 0-900 seconds following application of bradykinin (1 μM) alone or following pretreatment with the purinoceptor antagonist cocktail (Puri.: CGS 15943 (3 μM), RO4 (10 μM), and PPADS (100 μM)), D-AP5 (200 μM), or the purinoceptor antagonist cocktail and D-AP5 combined. One-way ANOVA (*N* = 5). **(g)** Change in nerve activity following application of bradykinin alone or following pretreatment with the purinoceptor antagonist cocktail. **(h)** Change in nerve activity following application of bradykinin alone or following pretreatment with D-AP5. **(i)** Change in nerve activity following application of bradykinin alone or following pretreatment with the purinoceptor antagonist cocktail and D-AP5. Bradykinin only dataset repeated in (g), (h), and (i) for ease of interpretation. (g-i) Two-way repeated-measures ANOVA (*N* = 5). GSK1016790A and bradykinin applied from t = 0-240 seconds. *\*P* > 0.05, \*\**P* > 0.01, \*\*\**P* < 0.001.

The contribution of ATP and glutamate to the activation of colonic afferents by GSK1016790A was evaluated using a cocktail of purinoceptor antagonists previously shown to block the afferent response to ATP (Figure S3), and the NMDA receptor antagonists D-AP5, kynurenic acid (KYNA), and L-701,324. The AUC of the afferent response to GSK1016790A (10 μM) was significantly reduced by the purinoceptor antagonist cocktail (*P* = 0.0263) and NMDA receptor antagonists D-AP5 (200 μM, *P* = 0.0041), KYNA (30 μM, *P* = 0.0263), and L-701,324 (100 nM, *P* = 0.0041, Figure 1e). Furthermore, combined incubation with both the purinoreceptor antagonist cocktail and D-AP5 virtually abolished afferent firing in response to GSK1016790A (*P* = 0.0006, Figure 1e). The inhibition of afferent firing to GSK1016790A by all pre-treatments persisted over time (*P* < 0.0001, Figure 1c-e, S4). By contrast, pre-treatment with either the purinoceptor antagonist cocktail, D-AP5 alone, or in combination had no significant effect on the AUC (*P* = 0.8367) and colonic afferent response to bradykinin (1 μM, *P* = 0.6189, Figure 1f-i). These findings indicate that ATP and glutamate are required for the activation of colonic afferents by GSK1016790A and that the respective antagonist pre-treatments are specific for the afferent response evoked by GSK1016790A rather than a generalized reduction in afferent sensitivity.

### 3.2 TRPV4-evoked colonic afferent activation requires the gut mucosa

Having established that ATP and glutamate are required for TRPV4-mediated stimulation of colonic afferents by GSK1016790A, we next sought to understand its site of action. Investigation of *Trpv4* expression alongside other algogenic receptors *Bdkrb2*, *Trpa1,* and *Trpv1* within our published scRNA-seq database of transcript expression in mouse colon-projecting neurons (Hockley et al., 2019, https://hockley.shinyapps.io/ColonicRNAseq/) revealed that expression of *Trpv4* is restricted to a small population of neurons (39 of 314 cells) in contrast to the wider expression of *Bdkrb2, Trpa1,* and *Trpv1* which are found in 206, 170, and 271 of the 314 cells studied, respectively (Figure 2a,b). The majority of cells expressing *Trpv4*, *Bdkrb2,* and *Trpa1* also co-expressed *Trpv1* (84.6%, 94.1%, and 94.7%, respectively), consistent with expression in nociceptors (albeit a small subpopulation for *Trpv4*) and role in nociception (Figure 2c-f). Given that GSK1016790A evokes a robust colonic afferent response greater than that evoked by bradykinin (Figure 1), in contrast to the restricted expression of *Trpv4* relative to *Bdkrb2* (or *Trpa1* and *Trpv1*) in colonic sensory neurons, we hypothesised that non-neuronal cell types contributed to the activation of colonic afferents by GSK1016790A. To interrogate this further, we examined the afferent response to GSK1016790A (10 μM) following removal of the colonic mucosa which abolished the AUC (*P* = 0.0123) and afferent response (*P* < 0.0001) to GSK1016790A (Fig. 2g,h). This was in marked contrast to the AUC and colonic afferent response to bradykinin (1 μM) or the TRPA1 agonist ASP7663 (100 μM) which were unchanged following removal of the colonic mucosa (AUC: *P* = 0.9537 and 0.5639, afferent response: *P* = 0.9927 and 0.5639, respectively, Figure 2i-l). These findings demonstrate that the mucosal expression of TRPV4 is required for the robust activation of colonic afferents by GSK1016790A.

**Figure 2.**
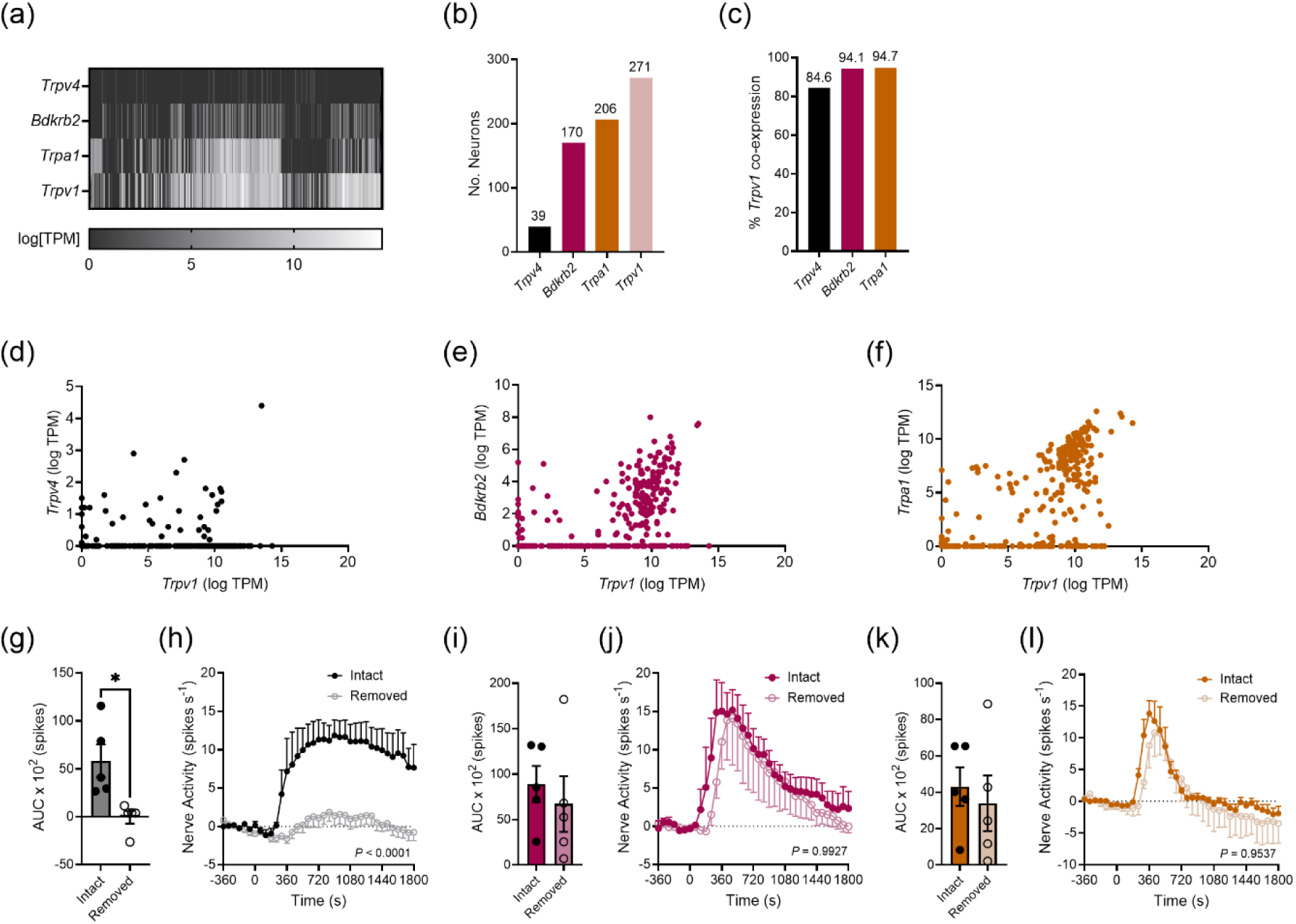
TRPV4 acts on the colonic mucosa to induce robust colonic afferent activity. **(a)** Heatmap of gene expression in murine colonic sensory neurons as log[TPM] (transcripts per million). **(b)** Number of neurons expressing *Trpv4, Bdkrb2,* and *Trpa1* transcripts in 314 cells tested. **(c)** % of *Trpv4-, Bdkrb2-,* and *Trpa1*-positive colonic sensory neurons co-expressing *Trpv1*. **(d)** Scatter plot of *Trpv1* co-expression with *Trpv4.* **(e)** Scatter plot of *Trpv1* co-expression with *Bdkrb2.* **(f)** Scatter plot of *Trpv1* co-expression with *Trpa1.* Each point denotes a single neuron. Data from (a-f) redrawn from Hockley *et al.,* 2019. **(g)** AUC of nerve activity from t = 0-900 seconds after the addition of GSK1016790A (10 μM) in preparations with the mucosa intact and removed. Unpaired *t* test (*N* = 5). **(h)** Change in nerve activity following application of GSK1016790A (10 μM) added from t = 0-240 seconds in preparations with the mucosa intact and removed. Two-way repeated-measures ANOVA (*N* = 5). **(i)** AUC of nerve activity from t = 0-900 seconds after the addition of bradykinin (1 μM) in preparations with the mucosa intact and removed. Unpaired *t* test (*N* = 5). **(j)** Change in nerve activity following application of bradykinin (1 μM) added from t = 0-240 seconds in preparations with the mucosa intact and removed. Two-way repeated-measures ANOVA (*N* = 5). **(k)** AUC of nerve activity from t = 0-900 seconds after the addition of ASP7663 (100 μM) in preparations with the mucosa intact and removed. Unpaired *t* test (*N* = 5). **(l)** Change in nerve activity following application of ASP7663 (100 μM) added from t = 0-240 seconds in preparations with the mucosa intact and removed. Two-way repeated-measures ANOVA (*N* = 5). \**P* < 0.05.

### 3.3 TRPV4 releases ATP to drive colonic afferent activation

To confirm that activation of TRPV4 by GSK1016790A evokes ATP release from the gut mucosa, we measured ATP levels in the basolateral compartment of human colon organoids stimulated apically with GSK1016790A, observing a significant increase in ATP release following addition of 1 μM of GSK1016790A with no sex-specific effect (Figure 3a,S5a). Consistent with this finding we also observed significant concentration dependent increases in ATP (Figure S5b) following incubation of male mouse colon with GSK1016790A, which were abolished at 10 μM by pre-treatment with HC067047 (10 μM, *P* < 0.0001, Figure 3b). Furthermore, pre-treatment with the NTPDase 1-3 inhibitor POM-1 (100 μM) produced a marked increase in the AUC (*P* = 0.0126) and colonic afferent response (*P* < 0.0001) to GSK1016790A (10 μM, Figure 3c-e) but had no effect on the AUC (*P* = 0.2021) and colonic afferent response to bradykinin (1 μM, *P* = 0.6781, Figure 3f-h), providing further evidence for a specific ATP-mediated increase in afferent activity following stimulation of TRPV4.

**Figure 3.**
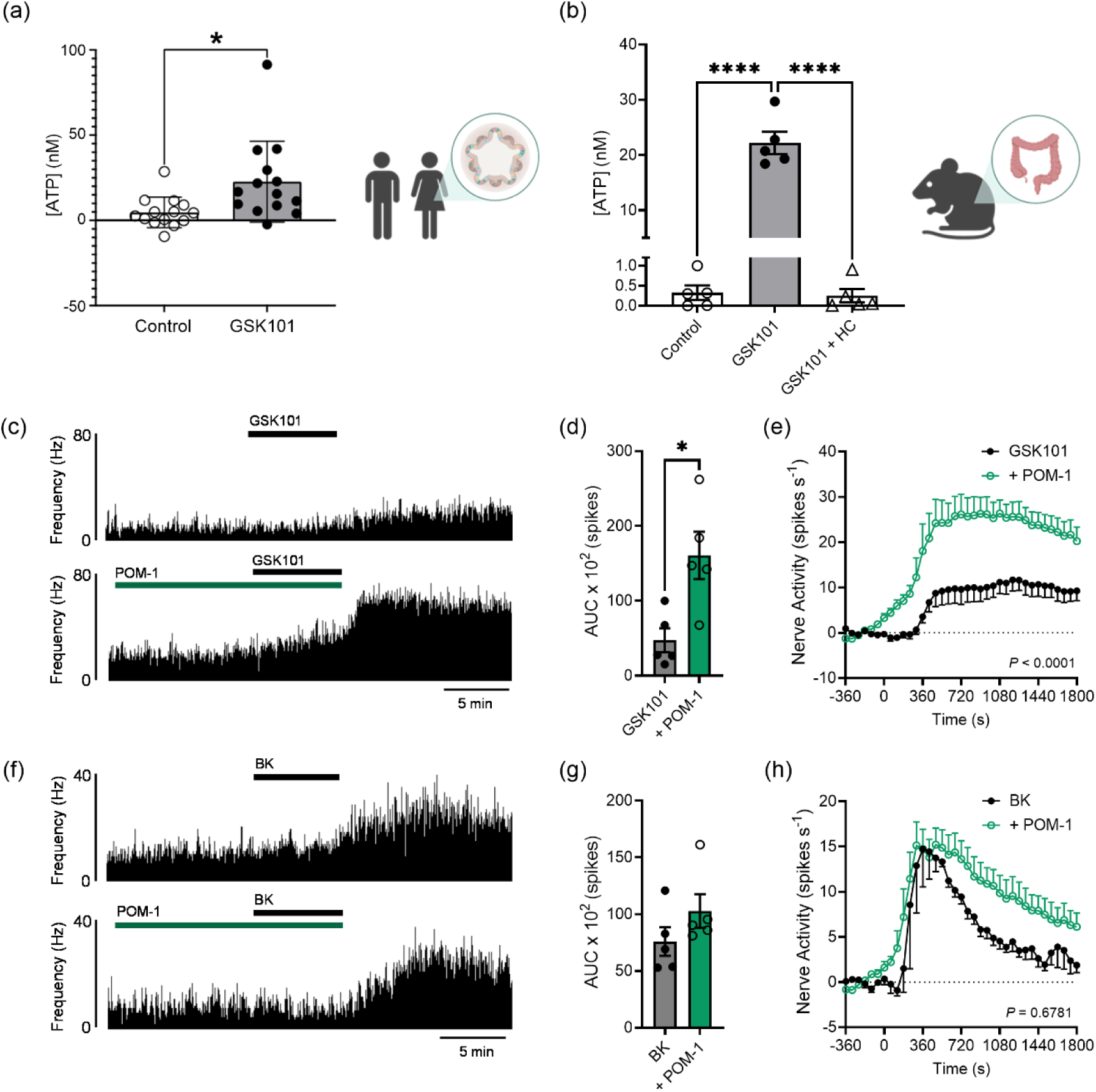
TRPV4-mediated colonic afferent activation requires ATP. **(a)** Differences in ATP concentration measured in the basolateral compartment of human colon organoids before and 10-minutes after apical exposure to DMSO (0.01%, control) or GSK1016790A (1 μM). Paired *t* test (*N* = 14). **(b)** ATP concentration measured from mouse colonic tissue supernatant after stimulation for 10-minutes with DMSO (0.01%, control), GSK1016790A (10 μM), or pretreatment for 10-minutes with HC067047 (HC, 10 μM) followed by GSK1016790A (10 μM) for 10-minutes. One-way ANOVA with Holm–Šídák’s multiple comparisons test (*N* = 5). **(c)** Rate histogram of LSN activity following application of GSK1016790A (10 μM) alone or following pretreatment with POM-1 (100 μM) in two separate nerve preparations. Bin width = 1-second **(d)** AUC of nerve activity from t = 0-900 seconds after application of GSK1016790A (10 μM) alone or following pretreatment with POM-1 (100 μM). Unpaired *t* test (*N* = 5). **(e)** Change in nerve activity after GSK1016790A (10 μM) added from t = 0-240 seconds alone or following pretreatment with POM-1 (100 μM). Two-way repeated-measures ANOVA (*N* = 5). (**f**) Rate histogram of LSN activity following application of bradykinin (1 μM) alone or following pretreatment with POM-1 (100 μM) in two separate nerve preparations. Bin width = 1-second. (**g**) AUC of nerve activity from t = 0-900 seconds after application of bradykinin (1 μM) alone or following pretreatment with POM-1 (100 μM). Unpaired *t* test (*N* = 5). **(h)** Change in nerve activity after bradykinin (1 μM) added at t = 0-seconds alone or following pretreatment with POM-1 (100 μM). Two-way repeated-measures ANOVA (*N* = 5). Illustrations produced using BioRender. \**P* < 0.05, \*\*\*\**P* < 0.0001.

### 3.4 TRPV4-driven colonic afferent activation is mediated by release of glutamate

As the subunits for AMPA, kainate, and excitatory metabotropic glutamate receptors are expressed alongside NMDA receptor subunits in colonic sensory afferents (Figure 4a), we next sought to confirm the contribution of glutamate and other excitatory glutamate receptors to the colonic afferent response to GSK1016790A. We detected a significant increase in glutamate release from the basolateral compartment of human colon organoids stimulated apically with GSK1016790A (1 μM, Figure 4b), with no sex-specific effect being observed (Figure S6a), establishing that GSK1016790A provokes glutamate release from the human intestinal mucosa. In addition, we also observed a significant increase in glutamate release from the mouse colon to 10 μM GSK1016790A (Figure S6b), which was abolished by pre-treatment with HC067047 (10 μM, *P* < 0.0001, Figure 4c) confirming that responses were TRPV4-mediated. Consistent with the release of glutamate and glutamate receptor expression in colonic neurons, the AUC of the afferent response to GSK1016790A (10 μM) was markedly reduced by pre-treatment with the combined AMPA/kainate blocker NBQX (10 μM, *P* = 0.0029) or the mGlu_5_ antagonist MTEP hydrochloride (100 μM, *P* < 0.0001, Figure 4d), effects which persisted over time (Figure 4e-f, *P* < 0.0001). These data demonstrate a role for AMPA/kainate and mGlu_5_ receptors alongside NMDA receptors in TRPV4-mediated colonic afferent activation.

**Figure 4.**
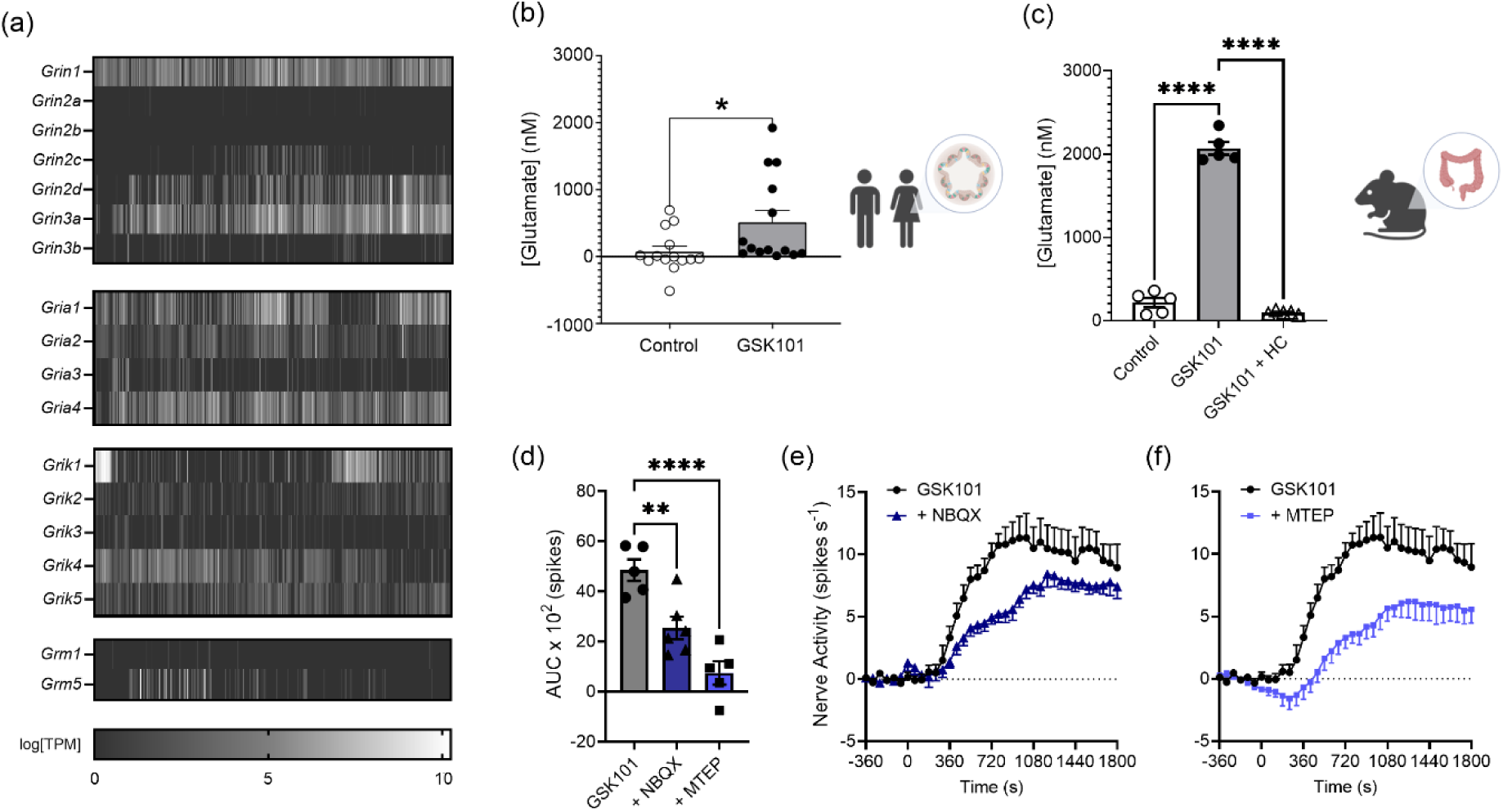
TRPV4-mediated colonic afferent activation requires glutamate. **(a)** Heatmap of gene expression in murine colonic sensory neurons as log[TPM] (transcripts per million) for subunits of NMDA, AMPA, and kainate receptors and excitatory metabotropic glutamate receptors. Data redrawn from Hockley *et al.,* 2019. **(b)** Differences in glutamate concentration measured in the basolateral compartment of human colon organoids before and 10-minutes after apical exposure to DMSO (0.01%, control) or GSK1016790A (1 μM). Paired *t* test (*N* = 14). **(c)** Glutamate concentration measured from mouse colonic tissue supernatant after stimulation for 10-minutes with DMSO (0.01%, control), GSK1016790A (10 μM), or pretreatment for 10-minutes with HC067047 (HC, 10 μM) followed by GSK1016790A (10 μM) for 10-minutes. One-way ANOVA with Holm–Šídák’s multiple comparisons test (*N* = 5). **(d)** AUC of nerve activity from t = 0-900 seconds following GSK1016790A alone or after pretreatment with NBQX (10 μM) or MTEP hydrochloride (100 μM). One-way ANOVA with Holm–Šídák’s multiple comparisons test compared to GSK1016790A alone (*N* = 5-6). **(e)** Change in nerve activity after GSK1016790A (10 μM) added from t = 0-240 seconds alone or following pretreatment with NBQX (10 μM). **(f)** Change in nerve activity after GSK1016790A (10 μM) added from t = 0-240 seconds alone or following pretreatment with MTEP hydrochloride (100 μM). GSK1016790A only dataset repeated in (e) and (f) for ease of comparison. (e-f) Two-way repeated-measures ANOVA (*N* = 5-6). Illustrations produced using BioRender. * *P* < 0.05, \*\**P* < 0.01, \*\*\*\**P* < 0.0001.

### 3.5 TRPV4 is expressed and functional in the gut mucosa

Finally, to confirm the contribution made by the mucosa to TRPV4-driven afferent activation, we examined the effect of GSK1016790A on intracellular Ca^2+^ concentration ([Ca^2+^]) in sensory neurons isolated from mouse DRG cultured alone or co-cultured with primary cells from the mouse colonic mucosa. Cells obtained from the mouse colonic mucosa predominantly comprised of epithelial cells and fibroblasts, and expressed transcripts corresponding to TRPV4 (Figure 5a-c). The application of GSK1016790A (100 nM and 1000 nM) to DRG neurons co-cultured with mucosal cells produced an increase in [Ca^2+^]_i_ of significantly greater magnitude compared to DRG neurons cultured alone (*P* = 0.0027, Figure 5d-f). Furthermore, GSK1016790A elicited an increase in [Ca^2+^]_i_ in a significantly greater proportion of co-cultured DRG neurons (*P* = 0.0025, Figure 5g), thus highlighting a difference in the mechanism of Ca^2+^ mobilisation and requirement for mucosal cells to fully realise the more marked response in DRG neurons to GSK1016790A. Comparison of the size of co-cultured DRG neurons demonstrated that smaller sized cells produced a response when stimulated by application of 1000 nM GSK1016790A indicative of the activation of a largely nociceptive population (*P* = 0.0062, Figure 5h). The increase in [Ca^2+^]_i_ within co-cultured DRG neurons to GSK1016790A (1000 nM) was significantly inhibited by pre-treatment with the TRPV4 antagonist HC067047 (10 μM*, P* = 0.0001), a purinoceptor antagonist cocktail (*P* = 0.0365), the NMDA receptor antagonist D-AP5 (*P* = 0.0417), and to an even greater extent by combining the purinoceptor antagonist cocktail and D-AP5 (*P* = 0.0038, Figure 5i). Overall, these results indicate that the DRG neuron response to GSK1016790A was mediated through the release of ATP and glutamate following activation of TRPV4.

**Figure 5.**
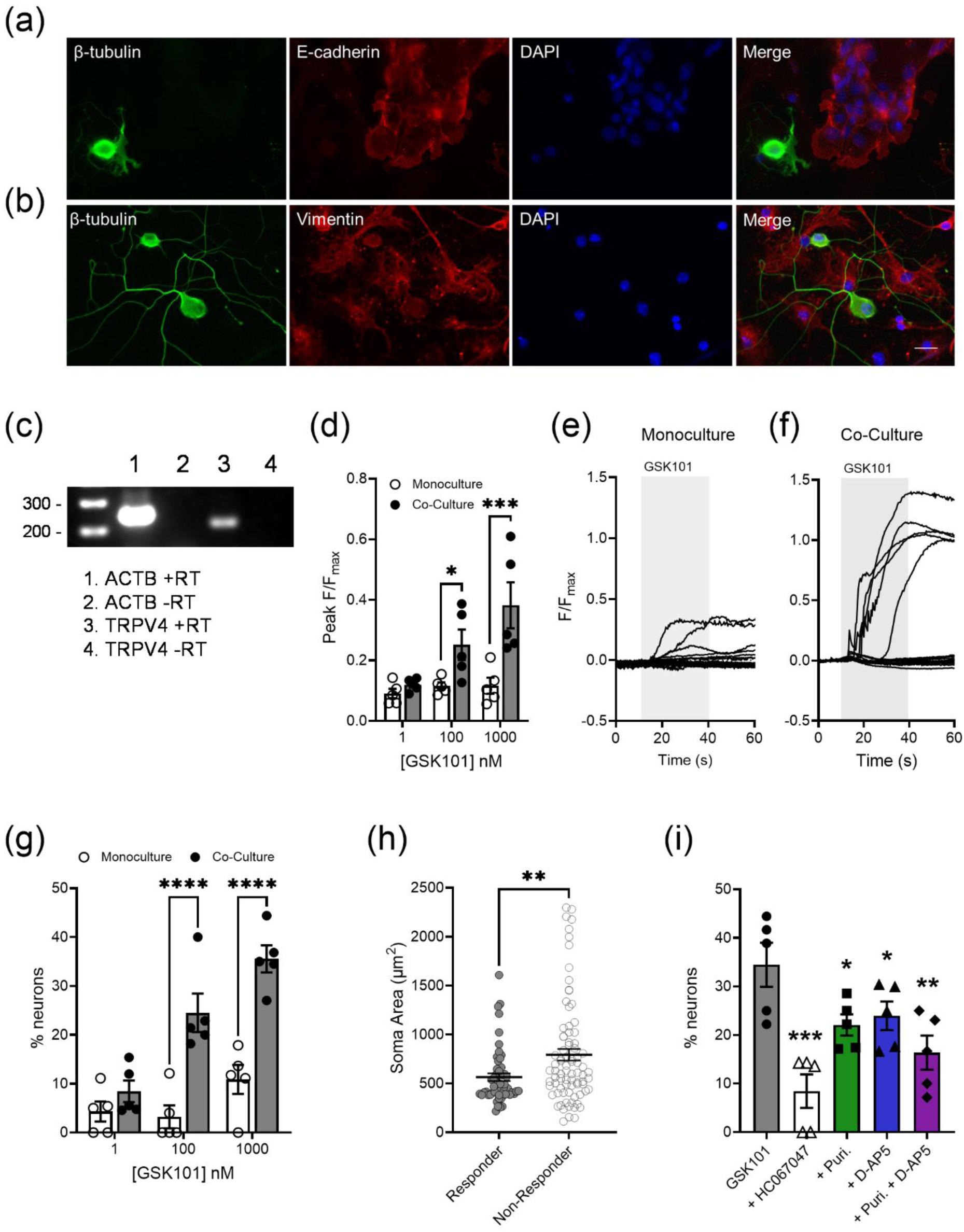
Mouse colonic mucosal cells enhance DRG neuron responses to GSK1016790A. **(a)** DRG neuron and colonic mucosal cell co-cultures stained with anti-β-tubulin antibody (green, neuronal marker), anti-e-cadherin antibody (red, epithelial marker), and DAPI (blue, nuclear marker). **(b)** DRG neuron and colonic mucosal cell co-cultures stained with anti-β-tubulin antibody (green), anti-vimentin antibody (red, fibroblast marker), and DAPI (blue). Scale bar = 20 μm. **(c)** RT-PCR analysis of TRPV4 and β-actin (ACTB) mRNA expression in primary cultured colonic mucosal cells. No signals were detected when reverse transcriptase (RT) was omitted. **(d)** Peak F/F_max_ values of GSK1016790A (1-1000 nM) responses from DRG neuron monocultures and co-cultured dishes. 100 nM: *P* = 0.0455, 1000 nM: *P* = 0.0002. Two-way ANOVA with Holm–Šídák’s multiple comparisons test (*n* = 5; *N* = 5, no. of dishes; no. of animals). **(e)** Representative response profiles from individual neurons in DRG neuron monocultures in response to GSK1016790A (1000 nM) applied from t = 10-40 seconds (shaded grey area). **(f)** Representative response profiles from individual neurons in co-cultures in response to GSK1016790A (1000 nM) applied from t = 10-40 seconds (shaded grey area). **(g)** Proportion of GSK1016790A (1-1000 nM) sensitive neurons in DRG neuron monocultures and co-cultured dishes. Two-way ANOVA with Holm–Šídák’s multiple comparisons test (*n* = 5; *N* = 5, no. of dishes; no. of animals). **(h)** Neuronal soma area for GSK1016790A (1000 nM) sensitive and insensitive DRG neurons from co-cultured dishes. Mann-Whitney U test (*N* = 54 (responder) and 81 (non-responder), no. of cells). **(i)** % of co-cultured DRG neurons responding to GSK1016790A (1000 nM) alone or following pretreatment with HC067047 (10 μM), the purinoceptor antagonist, D-AP5 (200 μM), and the purinoceptor antagonist cocktail and D-AP5 combined. One-way ANOVA with Holm–Šídák’s multiple comparisons test compared to GSK101670A alone (*n* = 5; *N* = 5, no. of dishes; no. of animals). \**P* < 0.05, \*\**P* < 0.01, \*\*\**P* < 0.001, \*\*\*\**P* < 0.0001.

## 4 Discussion

An improved understanding of how nociceptors are stimulated during gastrointestinal diseases is key to the development of new visceral analgesics. A large body of data has identified TRPV4 as an important effector of visceral pain and hypersensitivity making it an attractive therapeutic target for the treatment of gastrointestinal pain (Vergnolle, 2014). Recent single cell transcriptomic data indicates that only a small proportion of colonic-projecting neurons express TRPV4 suggesting that the pro-nociceptive effects of TRPV4 are also driven by non-neuronal cells (Hockley et al., 2019). Consequently, the goal of this study was to explore the contribution of the gut mucosa to TRPV4-mediated colonic afferent activation. Our findings revealed an essential role for the mucosa, which releases ATP and glutamate following agonist activation of TRPV4 to stimulate colonic afferents (Figure 6).

**Figure 6.**
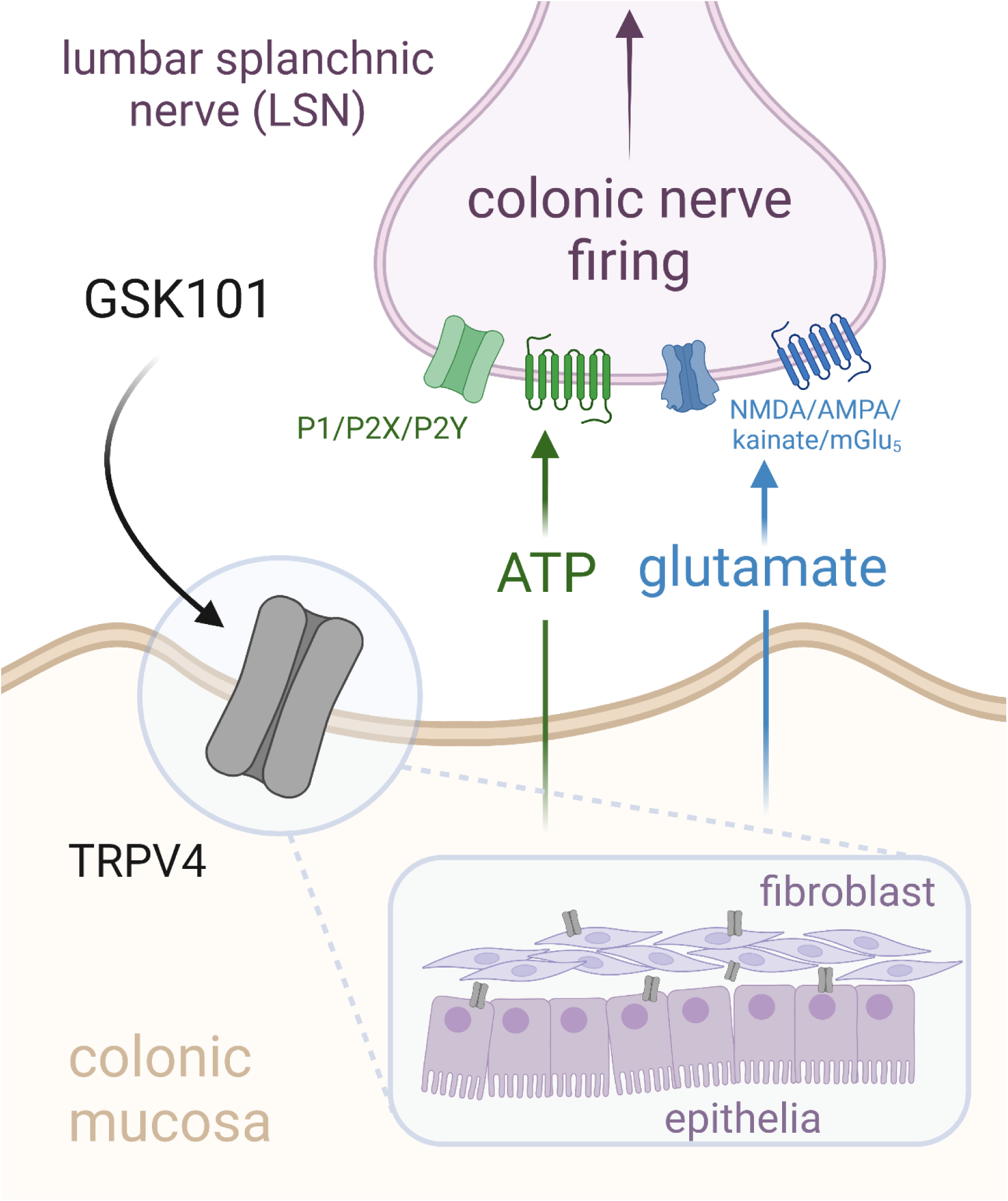
Proposed mechanism of colonic sensory nerve firing by TRPV4. GSK1016790A activates TRPV4 expressed upon cells of the colonic mucosa (most likely epithelial cells and fibroblasts). This drives the release of ATP and glutamate which act upon purinergic (P1, P2X, P2Y) and glutamatergic (NMDA, AMPA, kainate, mGlu_5_) receptors at the nerve terminal to drive LSN activation. Illustration produced using BioRender.

Within the study we observed a robust increase in colonic afferent firing in response to application of the TRPV4 agonist GSK1016790A which was abolished by pre-treatment with the TRPV4 antagonist HC067047, thus confirming selectivity of GSK1016790A for TRPV4. The stimulatory effect of GSK1016790A was lost following removal of the mucosa which demonstrates a mucosal site of action. Importantly, the colonic afferent response to both bradykinin and the TRPA1 agonist ASP7663 were unchanged between mucosa intact and removed preparations, demonstrating that removal of the mucosa had not damaged colonic innervation by nociceptors. Furthermore, the co-expression of TRPV4 with TRPA1 and the bradykinin B_2_ receptor in colonic sensory neurons would also indicate that removal of the mucosa had not affected the small population of TRPV4-expressing colonic neurons (Hockley et al., 2019).

The essential requirement for an intact gut mucosa to realise the afferent response to GSK1016790A suggests that the activation of TRPV4 promotes the release of mucosal mediators, which in turn stimulate colonic afferents. We hypothesized that this was driven by the release of ATP and glutamate, which we confirmed using several approaches. We demonstrated inhibition of the colonic afferent response to GSK1016790A in the presence of a pan-purinoreceptor antagonist cocktail and individual pre-treatments with NMDA, AMPA/kainate, or mGlu_5_ receptor antagonists, with combined pre-treatment with the pan-purinoreceptor antagonist cocktail and NMDA receptor antagonist D-AP5 abolishing the response to GSK1016790A. In addition, we measured increased ATP and glutamate release from the mouse colon following treatment with GSK1016790A which was repeated in primary human organoid cultures, the use of transwell organoid culture allowing confirmation of increased ATP and glutamate release from the basolateral compartment in response to apical GSK1016790A. Finally, we showed that marked activation of DRG neurons to GSK1016790A was only seen when co-cultured with primary colonic mucosa cells, an effect attenuated by individual and combined pre-treatment with a pan-purinoreceptor antagonist cocktail and D-AP5. A small proportion of DRG neurons responded to GSK1016790A when cultured alone, consistent with the restricted TRPV4 transcript expression in scRNA-seq databases (Hockley et al., 2019; Jung et al., 2023; Usoskin et al., 2015).

Our findings that TRPV4 stimulation promotes colonic afferent activation through mucosal ATP release is entirely consistent with previous work showing TRPV4-mediated release of ATP from epithelial cells in the oesophagus, stomach, lung, and bladder as well as intestinal cell lines (Aizawa et al., 2012; Bonvini et al., 2016; Mihara et al., 2011, 2016, 2018). Importantly, we demonstrate here that not only cell lines, but fully differentiated primary organoid cultures generated from surgically resected human bowel release ATP in response to TRPV4 agonist stimulation. This release was measured in the basolateral compartment, highlighting how ATP can be released in the vicinity of primary afferent endings. Studies in the lung and bladder also demonstrated a role for ATP in TRPV4-mediated sensory afferent firing (Aizawa et al., 2012; Bonvini et al., 2016), which we now confirm for the colon. In keeping with these and other studies looking at the colonic epithelium and cell lines (D’Aldebert et al., 2011; Liu et al., 2019), we found expression of TRPV4 in primary colonic mucosal cell cultures containing epithelial and fibroblast cells. While congruent with the published literature, we also showed that a cocktail of purinoreceptor antagonists targeting P1, P2X, and P2Y receptor subtypes is required to abolish the colonic afferent response to exogenous ATP, thereby highlighting the diversity of purinergic signalling in colonic afferents (Hockley et al., 2016; Shinoda et al., 2009).

Importantly, we revealed for the first time that TRPV4 activation promotes human intestinal epithelial cell glutamate release in keeping with i) the expression of glutamate transporters (excitatory amino acid transporter 3) on colonic epithelial cells (Holmseth et al., 2012; Hu et al., 2018), ii) the utilization of glutamate as a fuel source, and iii) expression of glutaminase in colonic enterocytes (Cherbuy et al., 1995), thereby allowing the synthesis of glutamate from glutamine. The released glutamate in turn stimulated colonic afferents through a combination of NMDA, AMPA/kainate, and mGlu_5_ receptor activation, consistent with the marked expression of these receptor subtypes in colon projecting sensory neurons (Hockley et al., 2019), supported by published work implicating NMDA and mGlu_5_ receptor activation in the development of visceral hypersensitivity (Lindström et al., 2008; McRoberts et al., 2001). Furthermore, given the significant contribution TRPV4 makes to colorectal afferent mechanosensitivity, our findings may also provide a mechanistic explanation for the inhibition of the VMR and pelvic afferent response to CRD following pre-treatment with glutamate receptor antagonists, namely TRPV4-mediated mucosal glutamate release (Brierley et al., 2008; Cenac et al., 2008, 2010) Similarly, our findings also suggest that TRPV4-mediated mucosal ATP release may contribute to the colonic afferent response to CRD in line with its subsequent inhibition by purinoreceptor antagonists, and studies showing that CRD promotes mucosal ATP release (Shinoda et al., 2009; Wynn et al., 2003). As such, further experiments are now warranted to confirm the role of TRPV4 in mechanically evoked ATP and glutamate release from the bowel and the contribution of specific epithelial and stromal cell types to TRPV4-evoked ATP and glutamate release.

Our findings that the majority of TRPV4-mediated colonic afferent activation is mucosally driven highlights the potential utility of mucosa restricted treatments for visceral pain in gastrointestinal disease. Indeed, many of the mediators implicated in the sensitisation of TRPV4 signalling in gastrointestinal disease also express receptors in gut epithelial cells and fibroblasts. The most pertinent of these is PAR2 which, like TRPV4, was not found to be widely expressed in colon projecting sensory neurons but is highly expressed in colonic epithelial cells (Green et al., 2000; Kong et al., 1997), its activation causing visceral hypersensitivity (Cenac et al., 2007; Vergnolle, 2004), and neurogenic inflammation (Nguyen et al., 2003), emphasising how the mucosa may also be a site of action for PAR2-mediated visceral hypersensitivity (Rolland-Fourcade et al., 2017), particularly through the modulation of TRPV4 signalling. Furthermore, significant increases in TRPV4 expression are found in mucosal biopsies from both people with irritable bowel syndrome and inflammatory bowel disease (Cheng et al., 2022; D’Aldebert et al., 2011; Fichna et al., 2015; Rizopoulos et al., 2018) suggesting that mucosal TRPV4 activation is likely to produce a greater effect on colonic afferent sensitisation in disease states. Given that mucosal TRPV4 expression has also been implicated in the development of colitis in people with inflammatory bowel disease (D’Aldebert et al., 2011), targeting mucosal TRPV4 activity may be effective for the treatment of both pain and inflammation in gastrointestinal diseases.

In summary, our study demonstrates that TRPV4-mediated colonic afferent activation is driven by the mucosal release of ATP and glutamate, which act on a broad array of glutamate and purinoreceptors to stimulate colonic afferents.

## 5 Additional Information

## Acknowledgements

This work was supported by the AstraZeneca Studentship (M. Y. Meng: G104109; L. W. Paine: G113502) and the UK Engineering and Physical Sciences Research Council (EPSRC) grant for the EPSRC Centre for Doctoral Training in Sensor Technologies for a Healthy and Sustainable Future (S. Oldroyd: EP/S023046/1).

---

**Supplementary Figure 1.**
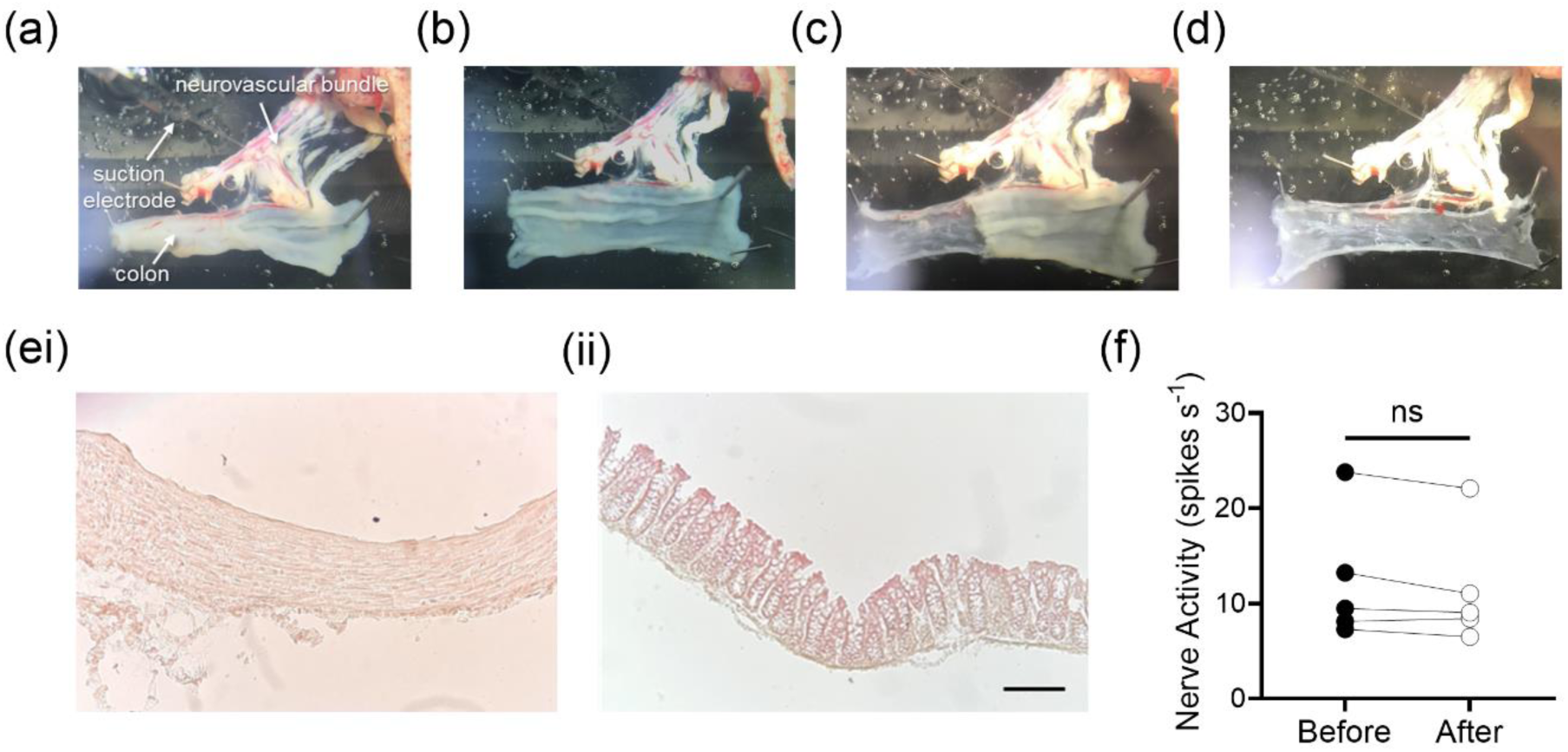
Removal of the colonic mucosa, validation of mucosa removal, and confirmation of tissue viability. **(a)** The colon is pinned directly to the base of the tissue bath and cut along the length of the mesenteric border. **(b)** The colon is pinned in each corner as a flat sheet with the mucosa facing upwards. **(c)** The top mucosal layer is carefully dissected free. **(d)** The underlying muscle is left intact. **(e)** Hemotoxylin and eosin staining of **(i)** the muscle layer remaining attached to the neurovascular bundle in the recording bath and **(ii)** the mucosal layer removed. Scale bar = 100 μm. **(f)** No significant difference was observed in the mean firing rate of baseline activity taken 6-minutes before and after the removal of the colonic mucosa (*P* = 0.1004, *N* = 5, paired *t* test).

**Supplementary Figure 2.**
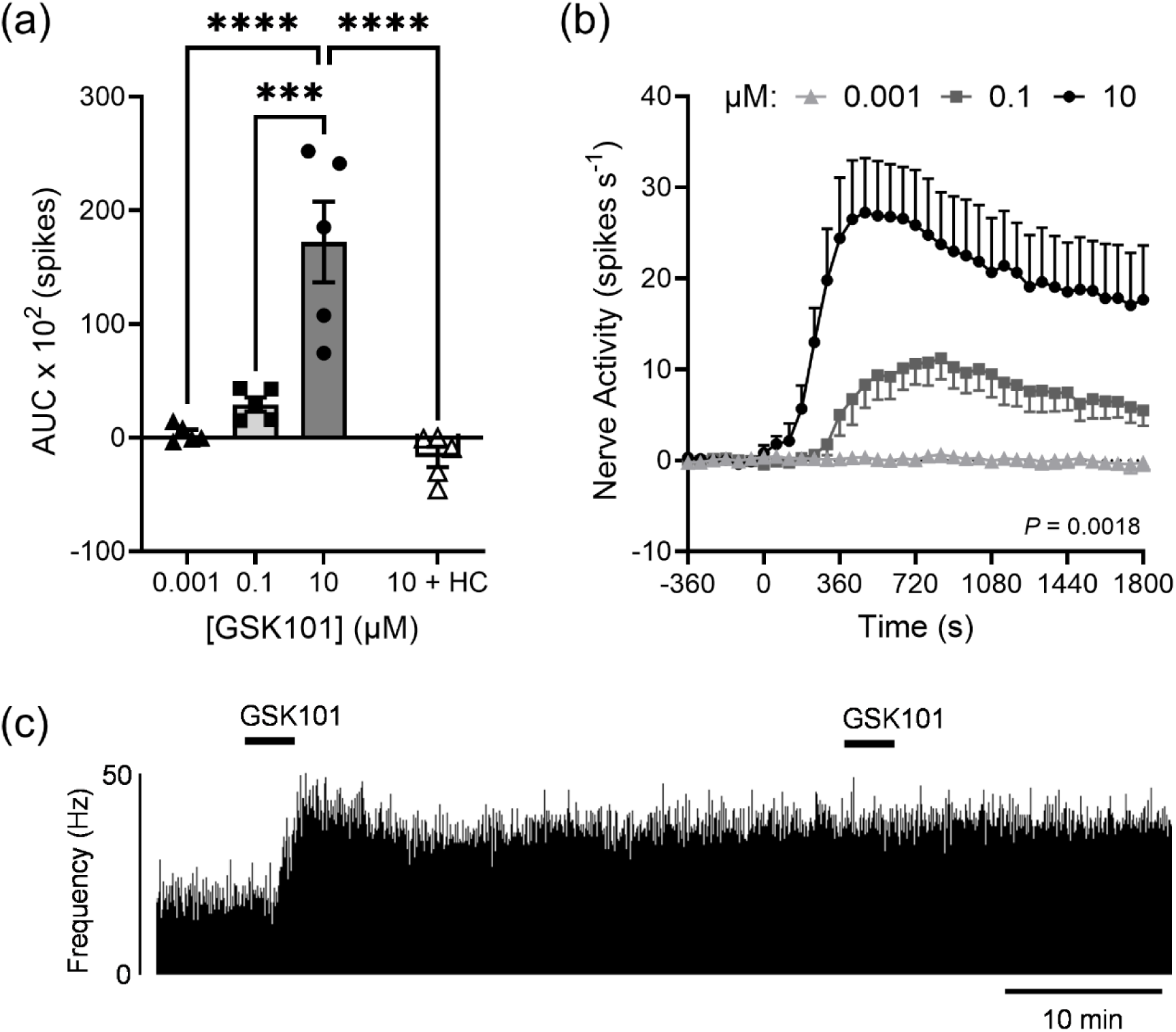
GSK1016790A evokes a dose-dependent increase in afferent firing which is desensitized upon repeat application. **(a)** AUC of nerve activity from t = 0-900 seconds following application of GSK1016790A (GSK101, 0.001-10 μM) or GSK1016790A (10 μM) following pretreatment with HC067047 (HC, 10 μM). One-way ANOVA with Holms-Šídák’s multiple comparisons test (*N* = 5). **(b)** Change in nerve activity after application of GSK1016790A (0.001-10 μM) from t = 0-240 seconds. Two-way repeated-measures ANOVA (*N* = 5). **(c)** Rate histogram of LSN activity following repeat application of GSK1016790A (10 μM) at an interval of 30 seconds. Bin width = 1 second. \*\*\**P* > 0.001, \*\*\*\**P* < 0.0001.

**Supplementary Figure 3.**
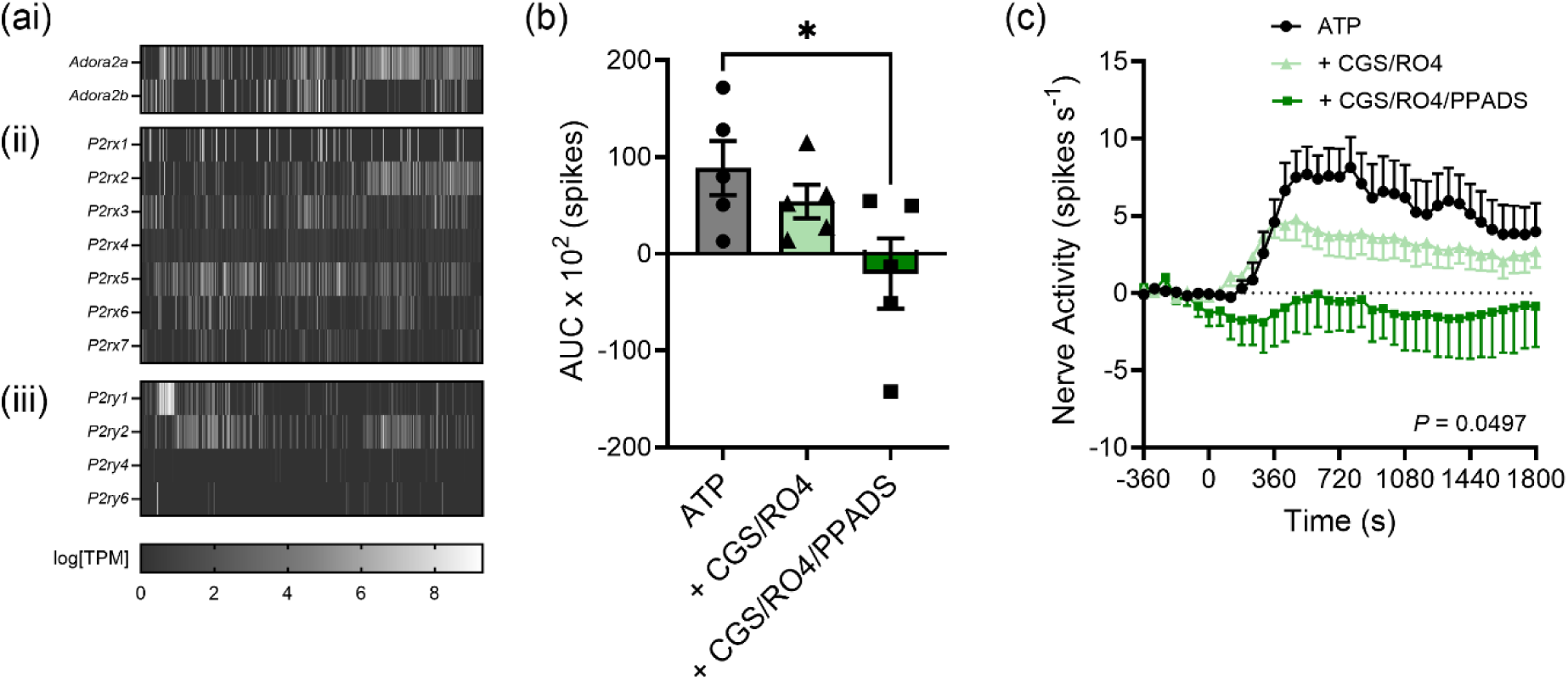
A combination of purinoceptor antagonists is required to inhibit the afferent response to ATP. **(a)** Heatmap of gene expression in murine colonic sensory neurons as log[TPM] (transcripts per million) for subunits which form excitatory subtypes of **(i)** P1, **(ii)** P2X, and **(iii)** P2Y purinergic receptors. Data redrawn from Hockley *et al.,* 2019. **(b)** AUC of nerve activity from t = 0-900 seconds following ATP (1 mM) alone or following pretreatment with CGS 15943 (3 μM) and RO4 (10 μM), or CGS, RO4, and PPADS (100 μM). ATP vs ATP + CGS/RO4/PPADS: *P* = 0.0190. One-way ANOVA with Holms-Šídák’s multiple comparisons test compared to ATP only (*N* = 5). **(c)** Change in nerve activity following application of ATP (1 mM) from t = 0-240 seconds alone or following pretreatment with CGS 15943 (3 μM) and RO4 (10 μM), or CGS 15943, RO4, and PPADS (100 μM). Two-way repeated-measures ANOVA (*N* = 5). \**P* < 0.05.

**Supplementary Figure 4.**
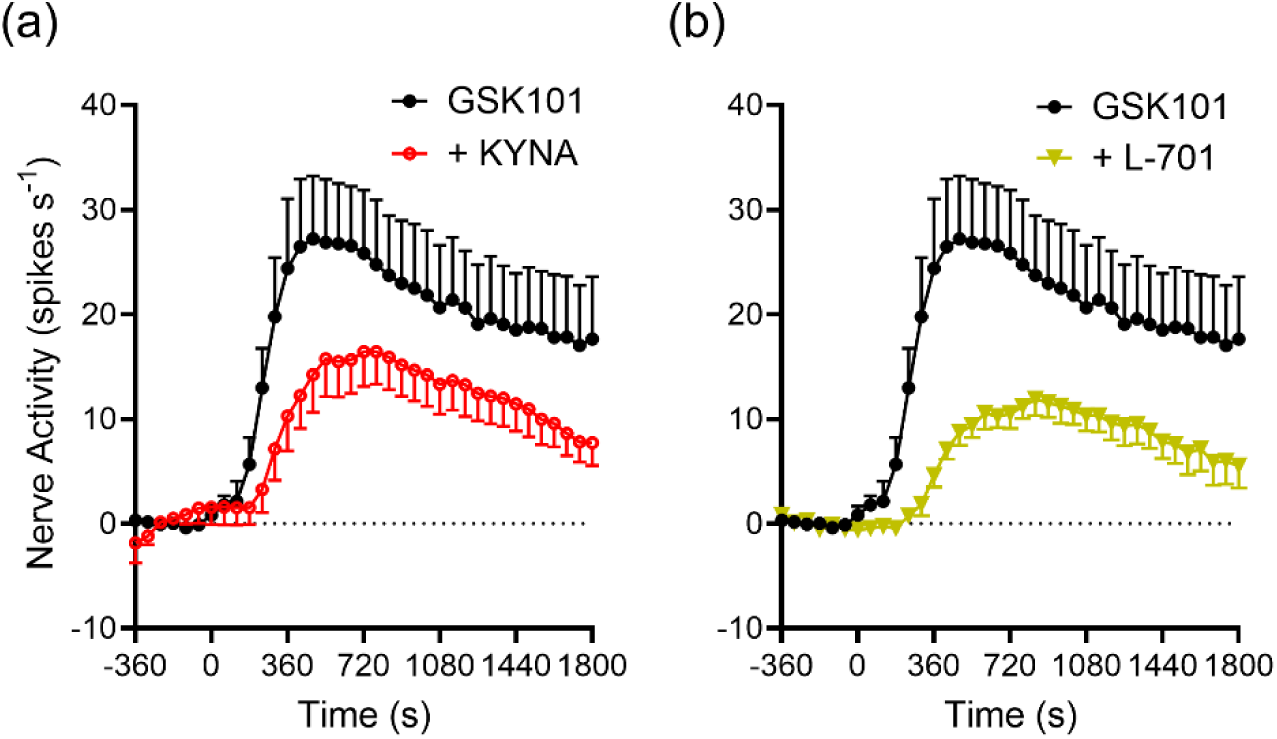
Inhibition of NMDA receptors reduces GSK1016790A-evoked colonic afferent activation. **(a)** Change in nerve activity following application of GSK1016790A (10 μM) from t = 0-240 seconds alone or following pretreatment with kynurenic acid (KYNA, 30 μM). **(b)** Change in nerve activity following application of GSK1016790A (10 μM) from t = 0-240 seconds alone or following pretreatment with L-701,324 (L-701, 100 nM). GSK1016790A only dataset repeated in (a) and (b) for ease of comparison. *P* < 0.0001, (a-b) Two-way repeated-measures ANOVA (*N* = 5-6).

**Supplementary Figure 5.**
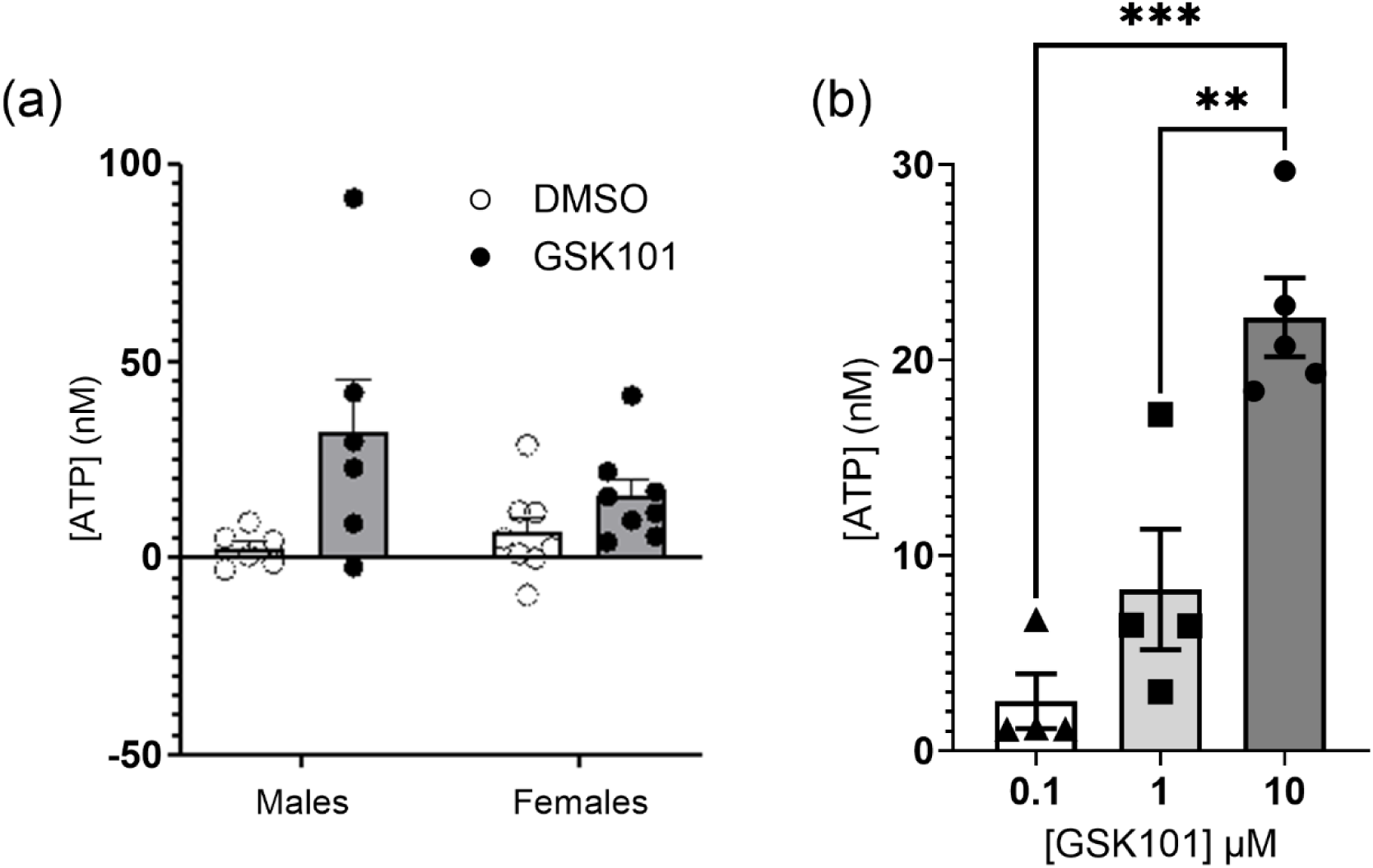
GSK1016790A evokes a sex-independent and dose-dependent increase in the release of ATP from human colon organoids and mouse colonic tissues respectively. **(a)** Differences in ATP concentration measured in the basolateral compartment of human colon organoids before and 10-minutes after apical exposure to DMSO (0.01%) or GSK1016790A (1 μM). Paired *t* test (*N* = 6 (males), *N* = 8 (females)). **(b)** Measurement of ATP release from mouse colonic tissue supernatant after stimulation for 10-minutes with GSK1016790A (0.1 - 10 μM). 0.1 vs 10: *P* = 0.0025, 1 vs 10: *P* = 0.0003. One-way ANOVA with Holms-Šídák’s multiple comparisons test (*N* = 4-5). \*\**P* < 0.01, \*\*\**P* < 0.001.

**Supplementary Figure 6.**
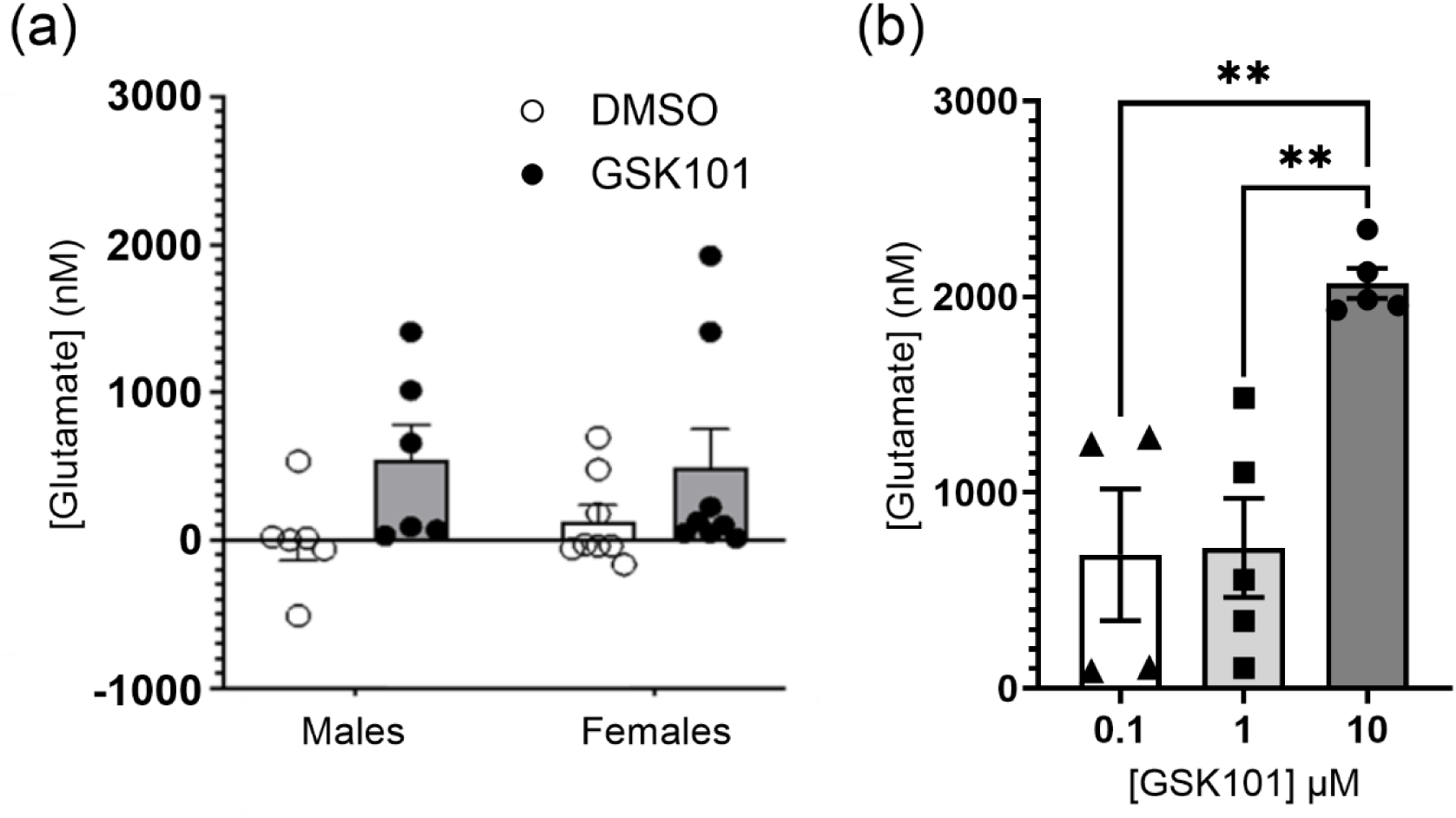
GSK1016790A evokes a sex-independent and dose-dependent increase in the release of glutamate from human colon organoids and from mouse colonic tissues respectively. **(a)** Differences in glutamate concentration measured in the basolateral compartment of human colon organoids before and 10-minutes after apical exposure to DMSO (0.01%) or GSK1016790A (1 μM). Paired *t* test (*N* = 6 (males), *N* = 8 (females)). **(b)** Measurement of glutamate release from mouse colonic tissue supernatant after stimulation for 10-minutes with GSK1016790A (0.1 - 10 μM). 0.1 vs 10: *P* = 0.0040, 1 vs 10: *P* = 0.0040. One-way ANOVA with Holms-Šídák’s multiple comparisons test (*N* = 4-5). \*\**P* < 0.01.

